# Thalamic and extra-thalamic connections of the Globus Pallidus in the human brain: The ultradirect pathway

**DOI:** 10.1101/688283

**Authors:** Zhong S. Zheng, Martin M. Monti

## Abstract

A dominant framework for understanding loss and recovery of consciousness, particularly in the context of severe brain injury, focuses on cortico-subcortical recurrent interactions, with a strong emphasis on excitatory thalamofugal projections. Recent work in healthy volunteers and patients, however, suggests a previously unappreciated role for the globus pallidus pars externa in maintaining a state of consciousness – a finding that is consistent with non-human animal work demonstrating the existence of direct (i.e., extrathalamic) pallido-cortical projections as well as their involvement in modulating electrocortical arousal and sleep. Leveraging on the high-quality Human Connectome Project dataset, we report for the first time in humans, in vivo evidence of (direct) pallido-cortical and pallido-thalamic projections, distinguishing between internal and external pallidal regions. Our data confirm, in humans, the existence of an “ultradirect” extra-thalamic pallido-cortical pathway, with the pars externa connecting preferentially, and extensively, to prefrontal cortex and the pars interna primarily connecting to sensorimotor cortical areas. Furthermore, we also report, for the first time in humans, the likely existence of a direct pathway uniting the globus pallidus pars externa and the medio-dorsal areas of thalamus often implicated in maintenance and recovery of consciousness. Consistent with the pallido-cortical connectivity results, the pars interna appeared to predominantly connect with the sensorimotor areas of thalamus. Collectively, these findings demonstrate the existence in humans of an extra-thalamic “ultradirect” pallido-cortical pathway and suggest a central role of the external segment of the globus pallidum in supporting consciousness.

## Introduction

The crucial role of cortico-subcortical interactions in the maintenance of waking consciousness has long been appreciated. Across studies in both healthy volunteers and patient populations, the function of these large-scale interactions has been shown to be modulated by the state of consciousness (Blumenfeld 2012, Kostopoulos 2001, Llinas et al 1998, McCafferty et al 2018, Monti 2012). While the content of consciousness, experience itself, might crucially rely on cortical sites, the thalamus is often considered to play a key role in allowing cortex to generate the type of neural activity that enables conscious experience (Alkire & Miller 2005), perhaps through enabling the delicate equilibrium of dynamic patterns of brain “cross-talk” which might underlie the emergence of specific aspect of information processing in the brain linking distant regions of the brain and/or enabling specific forms of information processing to emerge (Dehaene et al 2003, Demertzi et al 2019, Tononi 2008). Indeed, anesthesia-based loss of consciousness is well known to lead to altered activity in thalamic neurons (Andrada et al 2012), hypometabolism (Xie et al 2011) and decreased thalamo-cortical connectivity (Akeju et al 2014, Liu et al 2013), and a transition from tonic to a more rhythmic bursting firing mode (Silva et al 2010). Furthermore, thalamic stimulation, with many different approaches, has been shown to produce awakening (or faster awakening) from anesthesia (Alkire et al 2009, Alkire et al 2007, Yoo et al 2011) in the animal model. Conversely, thalamic lesions are believed to be associated with loss of consciousness (Castaigne et al 1981) and to be among the most prevalent signatures of protracted unconsciousness in post-mortem studies (Adams et al 2000, Adams et al 1999, Graham et al 2005). Moreover, both thalamic damage (Fernandez-Espejo et al 2011, Lutkenhoff et al 2015) and its de-afferentation from cortex (Lant et al 2016, Zheng et al 2017) have been demonstrated to be proportional, *in vivo*, to the depth of the impairment in patients with chronic disorders of consciousness (DOC).

Recent evidence, however, suggests that this thalamo-centric view underemphasizes the role of other structures within cortico-subcortical circuits in maintaining conscious wakefulness. Specifically, a growing number of studies involve the globus pallidus – in particular, its external segment (GPe) – in maintaining conscious wakefulness (Lazarus et al 2012, Qiu et al 2016b, Vetrivelan et al 2010). In the rodent model, optogenetic and deep brain stimulation (DBS) of this nucleus led to both increased sleep and EEG delta power (Qiu et al 2016a, Qiu et al 2016b) while cell-body specific lesion of GPe increased diurnal wake and decreased diurnal non-REM sleep (Qiu et al 2010). Moreover, the sleep promoting adenosine A2A receptors found densely in the striatum have been shown to innervate parvalbumin (PV) neurons of the rostral GPe (Yuan et al 2017). In humans, patients with Parkinson’s disease exhibit abnormal pallidal function (Bevan et al 2002, Gatev et al 2006, Hutchison et al 1994, Magnin et al 2000, Mallet et al 2008)) and commonly experience insomnia as part of non-motor symptomatology (Gjerstad et al 2007), with etiologies including sleep fragmentation, nocturnal immobility, REM sleep behavior disorder, among others (Chaudhuri et al 2006, Garcia-Borreguero et al 2003, Juri et al 2005, Partinen 1997, Trenkwalder 1998). In addition, findings from human consciousness research further demonstrate an association between the globus pallidus (though the external and internal segments were not differentiated) and level of arousal (Lutkenhoff et al 2015; Crone et al., 2017).

As the second largest component of the basal ganglia (BG), just after the striatum, GPe not only contains extensive connections with BG structures but also direct connections with frontal cortex (Chen et al 2015, Saunders et al 2015) and thalamus (Chattopadhyaya & Pal 2004, Hazrati & Parent 1991, Mastro et al 2014), both structures crucial for supporting consciousness. It has been proposed that, under the control of the dorsal striatum, direct GABAergic output from GPe to the frontal cortex (Chen et al 2015; Vetrivelan et al 2010) may be an important pathway for sleep-wake regulation (Qiu et al 2010; Yuan et al 2017) though the exact mechanism remains to be elucidated. If indeed direct connections between GPe and thalamus exist, this pathway could also contribute to the influence of cortical activity via the thalamo-cortical route. Thus far, direct GPe connections outside of the basal ganglia have only been verified in animals, but not yet in humans.

In this study, we aim to (i) test the existence in humans of “direct” GPe connections with cortex and thalamus – as assessed with *in vivo* methods, and (ii) contrast the pattern of connectivity of the GPe with that of the GPi, which has long been proposed to be part of the mesocircuit important for recovery of consciousness in DOC patients (Schiff 2010). To achieve these objectives, we employ high angular resolution diffusion imaging (HARDI) data from the Human Connectome Project (HCP; (Van Essen et al 2012)), which offer, as compared to conventional diffusion tensor imaging (DTI) approaches, the great advantage of resolving intra-voxel fiber heterogeneity, therefore providing higher spatial and angular resolution (Tuch et al 2002).

## Methods

### Data

We analyzed HARDI data provided by the HCP (Q3 Release), WU-Minn Consortium (http://www.humanconnectome.org) (Van Essen et al., 2012). 50 healthy subjects (26 females; 24 males; ages 22-35 years old) were included in the analysis. Imaging data were acquired on a modified 3T Siemens Skyra scanner. T1-weighted structural (TR = 2400 ms; TE = 2.14 ms; flip angle = 8°; FOV = 224×224 mm; voxel size = 0.7mm isotropic) and diffusion-weighted HARDI (TR = 5520 ms; TE = 89.5 ms; flip angle = 78°; FOV = 210×180 mm; voxel size = 1.25 mm isotropic; 3 shells of b = 1000, 2000, and 3000 s/mm^2^ with 90 diffusion directions per shell) data were used. Basic diffusion preprocessing steps have been applied which included B0 image intensity normalization, EPI distortions correction, eddy current correction, motion correction, gradient-nonlinearities correction, and registration of diffusion data with structural (Glasser et al., 2013). Preprocessing was accomplished using FSL tools (http://fsl.fmrib.ox.ac.uk).

### Regions of Interest (ROIs)

For cortical and thalamic ROIs, segmented FreeSurfer labels supplied by HCP were used (http://surfer.nmr.mgh.harvard.edu). To create the five distinct cortical zones (prefrontal [PFC], sensorimotor [SMC], posterior parietal [PPC], temporal [TEM], and occipital [OCC]), selected Desikan-Killiany atlas labels were combined together (See supplemental materials) (Desikan et al 2006). Since many of the BG ROIs were not available through FreeSurfer, GPe, GPi, striatum (STR), substantia nigra (SN), and subthalamic nucleus (STN) masks were obtained from the standard (Keuken & Forstmann 2015) probabilistic BG atlas and transformed into individual diffusion space. To refine the GPe and GPi masks, we used the individual whole pallidum FreeSurfer masks as a constraint to define the boundaries.

### Probabilistic Tractography

All imaging analyses were accomplished using FSL tools. First, the probability distribution function of fiber orientations at each voxel was estimated with the following parameters: 3 fibers per voxel, 1000 burn-ins, deconvolution model using zeppelins, and gradient nonlinearities considered. Next, probabilistic tractography was launched with 5000 samples drawn per voxel from each seed mask. For pallidocortical and pallidothalamic connectivity analysis, GPe and GPi each served as a seed mask with each of the five cortical targets and thalamus (THAL) as a waypoint mask, per hemisphere. A contralateral hemispheric mask was also included as an exclusion criterion to limit the connections ipsilaterally. This procedure generated ipsilateral pallidocortical and pallidothalalmic connections, though indirect connections arising from other nearby structures were also considered. For ease of referencing, we will refer to the results of this step as “indirect and direct” connections (IDC). To preclude the influence of indirect connections, we repeated probabilistic tractography with additional exclusion criteria, whereby streamlines entering any of the following masks were discarded: the remainder of BG (STR, SN, STN, and GPe or GPi) and cortical ROIs, thalamus, brainstem, cerebellum, as well as the contralateral hemisphere. The results from this analysis will be referred to as “direct connections” (DC) only. As a final analysis to more precisely visualize the different subregions within thalamus that GPe and GPi may be directly connected with, the thalamus also served as a seed with GPe and GPi as targets, following similar exclusion criteria. Next, the number of streamlines (i.e. connectivity strengths) that successfully reached the cortical and thalamic targets from the pallidal seeds were computed after applying a threshold of 50 (1% of 5000 samples per voxel) (Zhang et al 2014) to remove spurious connections for both IDC and DC. The results were then normalized and rescaled to account for individual differences in seed and target sizes (Eickhoff et al 2010). Individual streamline counts were divided by the total number of samples sent (5000 x seed mask size) and then rescaled by multiplying by the average of all total number of samples sent across all seeds and targets. This step accounted for differences in seed sizes. Next, the resulting values were then divided by the size of the targets and then again multiplied by the average size of all targets. This accounted for the variability in target sizes. For display of the tractograms, a threshold of 0.01% of the total number of streamlines sent was applied with an additional threshold of least 10% of subjects sharing the tracks. Segmentation of the thalamus based on differential pallidal connectivity was achieved through a winner-take-all approach on the normalized average group connectivity.

### Statistical Analysis

Repeated-measures ANOVAs of normalized and rescaled pallidal connectivity values were carried out for IDC and DC with seed (GPe, GPi), target (PFC, SMC, PPC, TEM, OCC, THAL), and hemisphere (left, right) as within-group factors. Upon finding significance, pairwise t-tests comparing GPe and GPi connections were performed along with multiple comparisons correction using the Benjamini & Hochberg (1995) method with false discovery rate set at q = 0.05. The new significance cutoff value was thus established at p < 0.02. Moreover, percent change of total streamline counts from IDC to DC was calculated.

## Results

### Assessing Pallidocortical and Pallidothalamic Connectivity Strengths

The total number of streamlines (normalized and rescaled) that successfully reached the cortical and thalamic targets from the pallidal seeds for both IDC and DC are plotted in Fig. 1. Statistical results are reported below:

**Figure 1.**
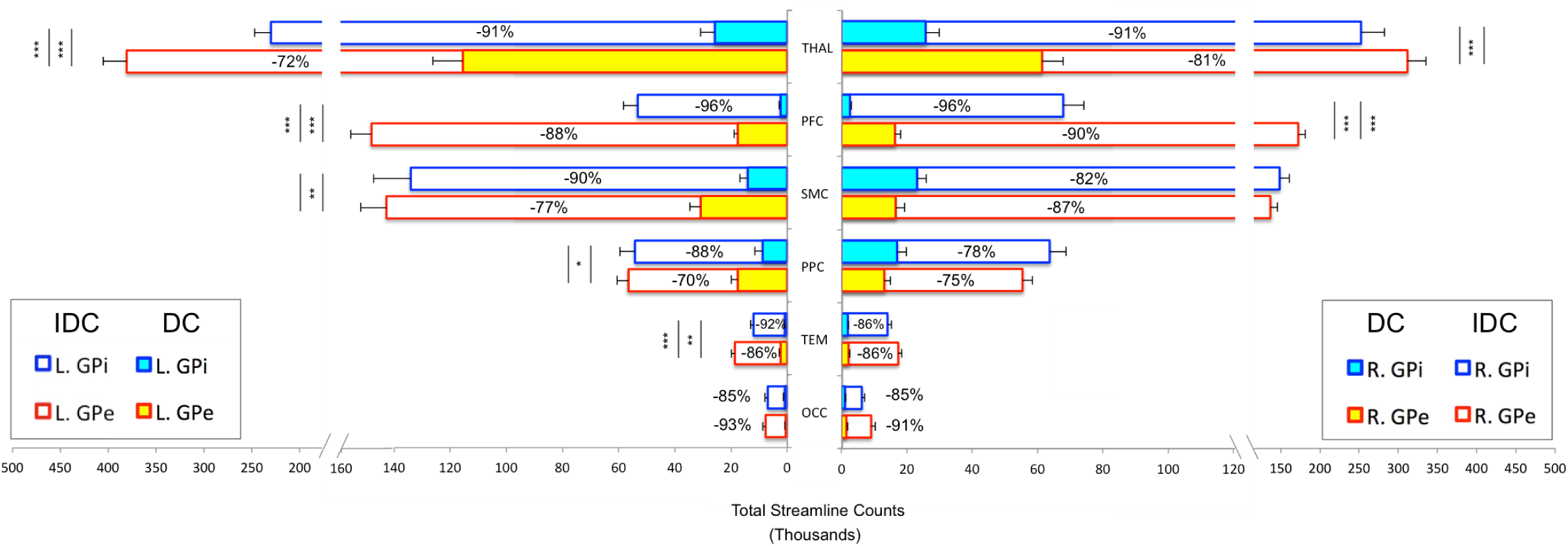
Pallido-cortical and pallido-thalamic connectivity strengths. Connectivity strengths as assessed by the group averages of total streamline counts between each seed and target ROI (normalized and rescaled) are plotted. DC (direct connections) and IDC (indirect and direct connections) are overlapped to reflect a percent decrease going from IDC to DC. GPe and GPi connections are statistically compared, with significance denoted using inner asterisks to correspond to DC and outer asterisks to IDC.

1. IDC: Significant main effects of seed (F = 109.063, p < 0.0001) and target (F = 153.47, p < 0.0001), as well as significant interactions of hemisphere × seed × target (F = 7.7, p < 0.01), hemisphere × seed (F = 11.2, p < 0.01), and seed × target (F = 46.9, p < 0.0001) were observed. Comparing GPe and GPi connectivity, post-hoc t-tests uncovered significantly higher connections for bilateral GPe-PFC, left GPe-THAL and GPe-TEM than GPi with these targets.

2. DC: Significant main effects of seed (F = 59.6, p < 0.0001), target (F = 106.5, p < 0.0001), and hemisphere (F = 15, p < 0.001), along with significant interactions of hemisphere × seed × target (F = 15.8, p < 0.0001), hemisphere × seed (F = 34.5, p < 0.0001), hemisphere × target (F = 18.9, p < 0.0001), and seed × target (F = 51.7, p < 0.0001) were detected. Post-hoc comparisons revealed significantly greater connections for bilateral GPe-PFC and GPe-THAL, left GPe-SMC, GPe-PPC, and GPe-TEM than GPi with these targets.

Overall, while the general pattern of DC remained roughly the same as IDC after introducing comprehensive exclusion criteria to minimize the influence of indirect connections, the direct connectivity values reduced drastically, with up to 96% decrease from IDC to DC, with GPi appearing to suffer more than GPe for the most of the connections (Fig. 1). We evaluated relative residual direct connectivity strength by separating the total streamline counts (tSC) into 3 categories: high (tSC > 10,000), medium (5,000 – 10,000 tSC), and low (tSC < 5,000) probability of connection. THAL, SMC, and PPC all exhibited medium to high probability of direct connections with GPe and GPi, but only GPe, not GPi, demonstrated a high likelihood of direct connection with PFC. Additional low direct connectivities were noted for pallidal connections with TEM and OCC regions.

### Topographical Organization of GPe and GPi Connections

Since our primary interest lies in direct connections, the descriptions hereafter will pertain only to DC. Group pallidal projections to the different cortical and thalamic targets as well as corresponding pallidal seed voxels with robust target connectivity are displayed in Fig. 2. Similar topographical connectivity patterns were identified for GPe and GPi, with the exception of PFC and THAL connections. While GPe-PFC connections covered the entire prefrontal target and originated from pallidal voxels concentrated mostly in the anterior subregion of GPe, GPi-PFC connections mainly projected to the posterior border of PFC (presumably, association motor cortices) and localized within the posterior subregion of GPi. Similar subregions of the GPe (anterior and a small posterior cluster) and GPi (posterior) corresponded to strong connections with THAL. Furthermore, SMC, PPC, TEM, and OCC connections resided primarily in posterior portions of GPe and GPi. Given the difficulty of dissociating preferential targeting in the thalamus from individual pallidothalamic tracks, connectivity-based segmentation of THAL with respect to GPe and GPi revealed distinct patterns of disparate pallidal organization within thalamus (Fig. 3). Namely, GPe connections occupied more medial aspects of thalamus, predominantly including putative midline, mediodorsal (MD), intralaminar (IL), and ventral anterior (VA) nuclei, with the highest concentration in the central medial intralaminar nucleus (CeM); GPi connections, on the other hand, were found in more lateral and posterior portions of thalamus, reflecting primarily ventral lateral (VL), ventral posterior lateral (VPL), ventral posterior medial (VPM), lateral posterior (LP), pulvinar (PUL), as well as some IL nuclei, with peak connections in VL motor thalamus. While both pallidal connections within THAL covered intralaminar nuclei, comparatively, GPe was more connected with central lateral (CL), central medial (CeM), and parafasicular (pf) nuclei, and GPi with centromedian (CM).

**Figure 2.**
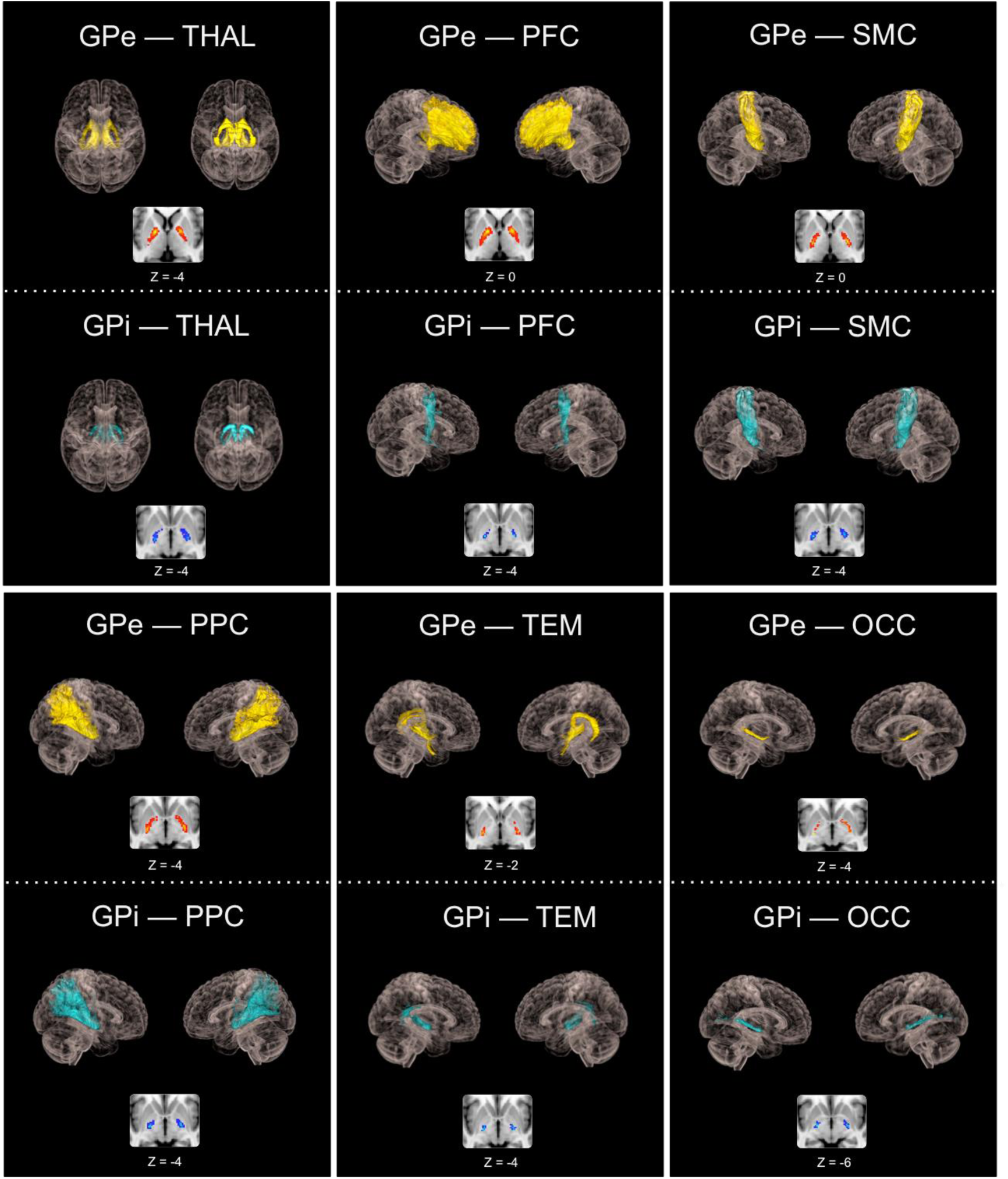
Topographical patterns of direct pallidal connectivity. Top sections show reconstructed pallidal pathways, and cropped axial images below reflect preferred subregional connectivity originating from GPe or GPi.

**Figure 3.**
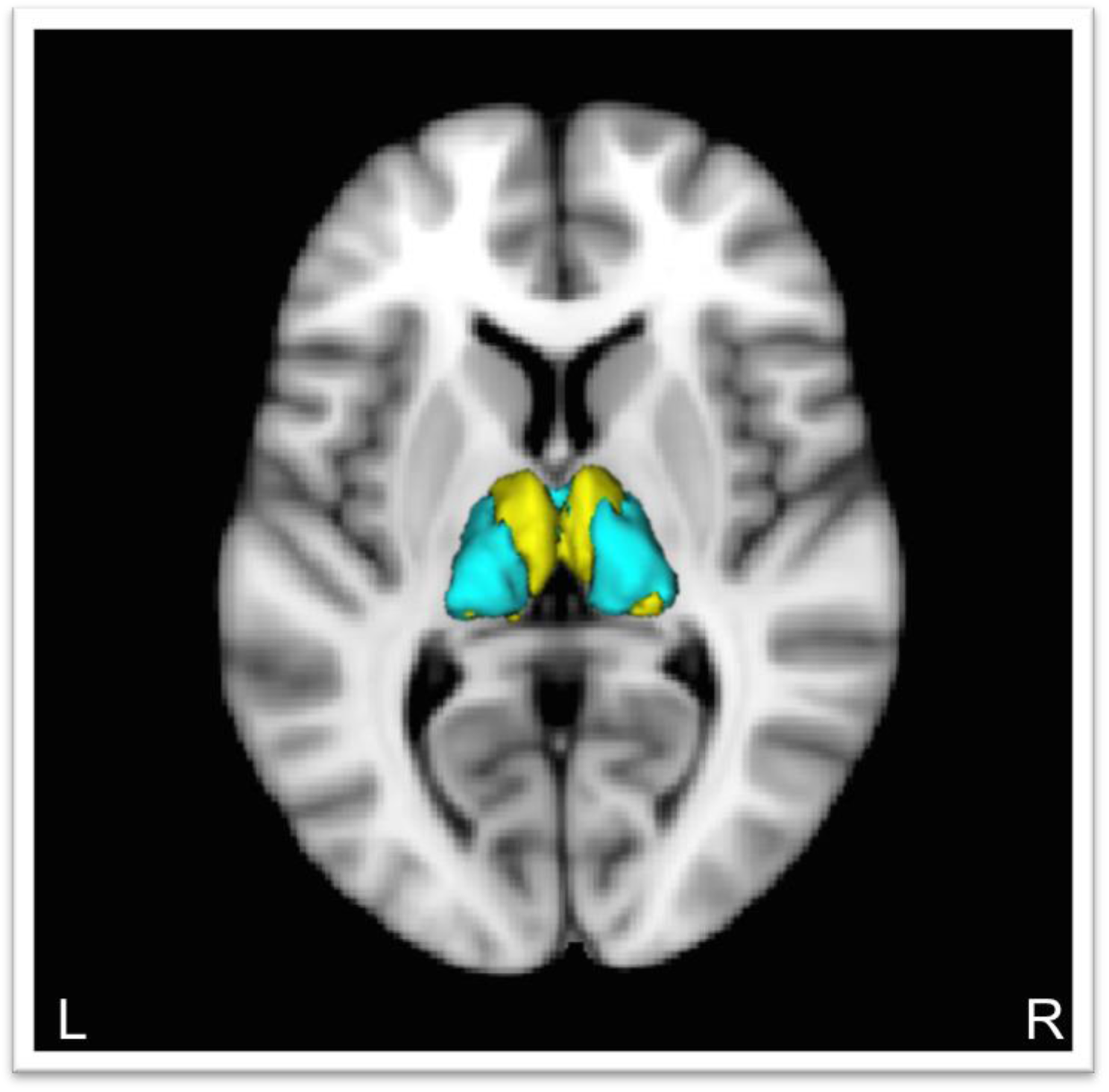
Segmentation of thalamus based on connectivity with GPe and GPi. Yellow = GPe connections within thalamus. Blue = GPi connections within thalamus

## Discussion

We used probabilistic tractography with comprehensive *a priori* exclusion criteria to characterize, for the very first time in humans, the pattern of direct pallidal connectivity with cortex and thalamus, separating its internal and external compartments. Overall, our findings of direct (i.e., extra-thalamic) connections between the GPe and prefrontal cortex and direct (i.e., not STN/GPi-mediated) connection to thalamus are not only consistent with animal studies (Chattopadhyaya & Pal, 2004; Chen et al 2015; Hazrati & Parent 1991; Mastro et al 2014; Saunders et al 2015), but also revealed novel findings including additional probable direct pallido-cortical connections and uncovering very different patterns of prefrontal and thalamic connectivity across the GPe and GPi. Indeed, the two compartments of the GP clearly mapped onto differential patterns of (direct) cortical and thalamic connectivity. A strong, widespread coverage of GPe tracks into the PFC, as compared to the weak GPi tracks, restricted to the posterior border of PFC (presumably, a partial association motor area). These pallido-PFC tracks also originated from different subregions of the pallidum, with anterior GPe corresponding to GPe-PFC projections, and posterior GPi representing GPi-PFC connections. Anterior GPe has been reported to connect with the prefrontal cortex (Francois et al 2004, Grabli et al 2004), whereas the posterior portion of GPi has been well established as a sensorimotor region (Visser-Vandewalle et al 2009). While the strong, direct GPe-PFC would still survive a more stringent thresholding, the weak, direct GPi-PFC connectivity would most likely fail. The direct GPe-PFC connections are validated against animal tracer studies and are thus likely to exist, but direct GPi-PFC connections may be more limited and if present, perhaps, exclusive to sensorimotor regions. Of course, given the structural nature of our methodology, the role of the direct GPe-PFC pathway, in humans, remains speculative. Findings from animal studies suggest that GPe may be involved in regulating sleep-wake (SW) behavior and cortical activity (Qiu et al 2016a, Qiu et al 2010, Qiu et al 2016b, Vetrivelan et al 2010). GPe lesions in rodents produced a 45% increase in wakefulness and significant slowing of cortical EEG (Qiu et al 2010), while DBS and optogenetic excitation of GPe neurons have been shown to directly promote sleep (Qiu et al 2016a, Yuan et al 2017). Confirming a separation between the duties of the two compartments of the pallidum, lesions to the GPi did not significantly change SW activity (Qiu et al 2010).

In addition to the direct pathway from GPe to frontal cortex, a direct connection between GPe and thalamus may also have important implications in disorders of consciousness. The mesocircuit hypothesis of DOC posits that following structural damages, functional disruptions at the BG-THAL level may also occur, leading to a reduced excitation of the anterior forebrain (Schiff 2010). Thalamo-prefrontal connections have been advocated to be necessary for sustaining organized behavior during wakefulness (Schiff 2008) and consistently implicated in DOC (Laureys et al 2000, Monti et al 2015, Zheng et al 2017). Yet, the current hypothesis fails to factor GPe into the model and attributes the reduced cortical excitation to the undertaking of the GPi through its excessive inhibition supposedly on the central thalamic nuclei (intralaminar complex and adjacent paralaminar portion of association nuclei—MD, VA, VL, and PUL) following insufficient inhibition from the striatum. Nonetheless, we could not find a study to date presenting evidence indicating that GPi may be the pallidal structure responsible for the down-regulation of frontal activity. In fact, due to the difficulty of separating the pallidum into internal and external parts from limited resolution, studies investigating DOC have only reported the globus pallidus as whole to be involved. For example, DOC patients, compared to controls, showed reduced metabolisms in the striatum and central thalamus yet increased metabolism in the globus pallidus (Fridman et al 2014). Moreover, behavioral arousal in chronic DOC patients has been shown to be negatively correlated with degree of atrophy in the dorsal striatum and globus pallidus (Lutkenhoff et al 2015). In the anesthesia model, once direct pallido-cortical connections were included in the directed connectivity modeling, propofol-induced loss of consciousness was found to disrupt pallido-cortical, but not thalamo-cortical connectivity (Crone et al 2017). These observations emphasize a critical role of the globus pallidus in supporting consciousness and do not preclude the possibility that GPe may be a vital contributor.

Nevertheless, our findings offer support for GPe as a potential influencer in the mesocircuitry. The direct connection between GPe and PFC, as shown in our results and prior animal studies, may serve to suppress neurons in the frontal cortex, though it is unclear whether the targets are pyramidal neurons or interneurons, or both. Another indication of GPe being intimately involved with PFC is evident in the pallidal connectivity based segmentation of the thalamus, where GPe displayed a greater preference for medial thalamus, which contains the main prefrontal projecting thalamic nucleus—MD (Klein et al 2010) and also a part of the central thalamus. Impairment in the mediodorsal nucleus stands as one of the most replicated findings in the DOC literature (Fernandez-Espejo et al 2010, Hannawi et al 2015, Lutkenhoff et al 2015, Lutkenhoff et al 2013, Monti et al 2015, Zheng et al 2017). Furthermore, the strongest thalamic connection with the GPe was detected in the central medial nucleus, a component of the anterior intralaminar complex. Receiving inputs from the brainstem and basal forebrain, the anterior intralaminar and mediodorsal thalamus have been postulated to assume a role in arousal regulation and possibly extending to awareness (Schiff 2008). On the other hand, GPi was most connected with ventral lateral nucleus, implying a more motor-related function, consistent with the classic role of GPi. This is additionally reflected in GPi’s projections to more motor parts of the frontal cortex.

Administration of zolpidem, a selective GABA-ω1 (a subunit of GABA-A receptor complex) agonist commonly used to treat insomnia, has produced paradoxical effects in DOC patients that have led to increased arousal and cognitive performance (Chatelle et al 2014, Clauss & Nel 2006, Clauss et al 2000, Shames & Ring 2008, Whyte et al 2014). Although the mechanism of action of this paradoxical response is largely unknown, the notion that zolpidem could act on the GABA-A receptors found abundantly in the globus pallidus to shut down its inhibitory output has been considered. Seeing that both GPe and GPi may contain large numbers of GABA-A receptors (Xue et al 2010), and that differing patterns of frontal and thalamic connectivity exist between the two pallidal structures, we propose that zolpidem’s action on GPi may, if detectable, improve more motor-related aspects of behavior, whereas zolpidem’s target on GPe may underlie the actual arousal and possibly cognitive increase observed in the DOC patients as GPe has been implicated in arousal regulation and is robustly connected with higher-order cognitive areas. Additional evidence alluding to a greater likelihood of GPe than GPi in contributing to the positive effects of zolpidem stems from findings that collectively demonstrate changes in prefrontal activity following administration of zolpidem in DOC patients who responded to the medication (Brefel-Courbon et al 2007, Chatelle et al 2014, Clauss et al 2000, Williams et al 2013). Again, GPe’s preferred connections with the PFC directly, and indirectly through the medial thalamus, strengthen our hypothesis. An update to the mesocircuit hypothesis with inclusion of the GPe is thus needed.

We found additional probable direct connections for both pallidal structures with SMC and PPC, although there currently lacks evidence from animal studies to validate these findings. These pallido-cortical connections originated from the posterior portions of the pallidum, which is consistent with DTI findings in humans from Draganski et al (2008), but the authors did not use exclusion criteria to limit indirect connections. However, while quantitatively different, DC (direct connections) versus IDC (indirect + direct connections) may remain qualitatively similar, sharing the same topographical patterns of connectivity.

Using diffusion imaging to infer the existence of connections between regions presents many challenges, especially for determining “direct” connections. Currently, one of the only ways of approaching the delineation of “direct” connections relies on strategically implementing exclusion criteria based on some a priori knowledge. However, the selection of exclusion criteria stands to be difficult and presents a tradeoff. On the one hand, not adding enough exclusions may still be subjected to the influence of indirect connections, but on the other hand, including too many exclusions may be overly stringent and lead to false negatives. Because our primary goal was to demonstrate the existence of “direct” connections, we opted for extensive exclusion criteria, albeit overly stringent, in hopes of removing as much of the indirect connections as possible. Though there could still exist some indirect connections in our “direct” connectivity results, the majority of which are likely addressed. On a general note, the inability to differentiate afferent and efferent connections remains a major limitation of diffusion tractography, although animal tracer studies suggest an output pathway from GPe to PFC and potentially bi-directional connections between GPe and thalamus.

## Conclusions

We demonstrated with probabilistic tractography using HARDI data provided by the HCP that direct GPe connections with PFC and THAL are likely to exist in humans, concurrent with animal tracer studies. These direct GPe connections also exhibited differing patterns of frontal and thalamic connectivity when compared against GPi. Favoring connections with distributed prefrontal cortex and medial thalamus, GPe is situated in a position to influence key aspects of consciousness. Conversely, GPi, preferring connections with more motor-related regions, remains a central player in the regulation of motor control. The current findings urge for an update to the mesocircuit hypothesis with the incorporation of GPe to shed light on the mechanisms underlying disorders of consciousness.

## Acknowlegments

This work was in part funded by the Tiny Blue Dot foundation. Data were provided by the Human Connectome Project, WU-Minn Consortium (Principal Investigators: David Van Essen and Kamil Ugurbil; 1U54MH091657) funded by the 16 NIH Institutes and Centers that support the NIH Blueprint for Neuroscience Research; and by the McDonnell Center for Systems Neuroscience at Washington University.

## References

Adams JH, Graham DI, Jennett B. 2000. The neuropathology of the vegetative state after an acute brain insult. Brain 123 (Pt 7): 1327–3.

Adams JH, Jennett B, McLellan DR, Murray LS, Graham DI. 1999. The neuropathology of the vegetative state after head injury. J Clin Pathol 52: 804–6

Akeju O, Loggia ML, Catana C, Pavone KJ, Vazquez R, et al. 2014. Disruption of thalamic functional connectivity is a neural correlate of dexmedetomidine-induced unconsciousness. Elife 3: e04499

Alkire MT, Asher CD, Franciscus AM, Hahn EL. 2009. Thalamic microinfusion of antibody to a voltage-gated potassium channel restores consciousness during anesthesia. Anesthesiology 110: 766–7.

Alkire MT, McReynolds JR, Hahn EL, Trivedi AN. 2007. Thalamic microinjection of nicotine reverses sevoflurane-induced loss of righting reflex in the rat. Anesthesiology 107: 264–7.

Alkire MT, Miller J. 2005. General anesthesia and the neural correlates of consciousness. Prog Brain Res 150: 229–4.

Andrada J, Livingston P, Lee BJ, Antognini J. 2012. Propofol and etomidate depress cortical, thalamic, and reticular formation neurons during anesthetic-induced unconsciousness. Anesth Analg 114: 661–9

Bevan MD, Magill PJ, Terman D, Bolam JP, Wilson CJ. 2002. Move to the rhythm: oscillations in the subthalamic nucleus-external globus pallidus network. Trends Neurosci 25: 525–3.

Blumenfeld H. 2012. Impaired consciousness in epilepsy. Lancet Neurol 11: 814–2.

Brefel-Courbon C, Payoux P, Ory F, Sommet A, Slaoui T, et al. 2007. Clinical and imaging evidence of zolpidem effect in hypoxic encephalopathy. Ann Neurol 62: 102–5

Castaigne P, Lhermitte F, Buge A, Escourolle R, Hauw JJ, Lyon-Caen O. 1981. Paramedian thalamic and midbrain infarct: clinical and neuropathological study. Ann Neurol 10: 127–4.

Chatelle C, Thibaut A, Gosseries O, Bruno MA, Demertzi A, et al. 2014. Changes in cerebral metabolism in patients with a minimally conscious state responding to zolpidem. Front Hum Neurosci 8: 917

Chattopadhyaya R, Pal A. 2004. Improved model of a LexA repressor dimer bound to recA operator. J Biomol Struct Dyn 21: 681–9

Chaudhuri KR, Healy DG, Schapira AH, National Institute for Clinical E. 2006. Non-motor symptoms of Parkinson’s disease: diagnosis and management. Lancet Neurol 5: 235–4.

Chen MC, Ferrari L, Sacchet MD, Foland-Ross LC, Qiu MH, et al. 2015. Identification of a direct GABAergic pallidocortical pathway in rodents. Eur J Neurosci 41: 748–5.

Clauss R, Nel W. 2006. Drug induced arousal from the permanent vegetative state. NeuroRehabilitation 21: 23–8

Clauss RP, Guldenpfennig WM, Nel HW, Sathekge MM, Venkannagari RR. 2000. Extraordinary arousal from semi-comatose state on zolpidem. A case report. S Afr Med J 90: 68–7.

Crone JS, Lutkenhoff ES, Bio BJ, Laureys S, Monti MM. 2016. Testing Proposed Neuronal Models of Effective Connectivity Within the Cortico-basal Ganglia-thalamo-cortical Loop During Loss of Consciousness. Cerebral cortex (New York, N.Y. : 1991)

Dehaene S, Sergent C, Changeux JP. 2003. A neuronal network model linking subjective reports and objective physiological data during conscious perception. Proc Natl Acad Sci U S A 100: 8520–5

Demertzi A, Tagliazucchi E, Dehaene S, Deco G, Barttfeld P, et al. 2019. Human consciousness is supported by dynamic complex patterns of brain signal coordination. Science 5: eaat7603

Draganski B, Kherif F, Kloppel S, Cook PA, Alexander DC, et al. 2008. Evidence for segregated and integrative connectivity patterns in the human Basal Ganglia. J Neurosci 28: 7143–52.

Fernandez-Espejo D, Bekinschtein T, Monti MM, Pickard JD, Junque C, et al. 2011. Diffusion weighted imaging distinguishes the vegetative state from the minimally conscious state. Neuroimage 54: 103–1.

Fernandez-Espejo D, Junque C, Bernabeu M, Roig-Rovira T, Vendrell P, Mercader JM. 2010. Reductions of thalamic volume and regional shape changes in the vegetative and the minimally conscious states. J Neurotrauma 27: 1187–9.

Francois C, Grabli D, McCairn K, Jan C, Karachi C, et al. 2004. Behavioural disorders induced by external globus pallidus dysfunction in primates II. Anatomical study. Brain 127: 2055–7.

Fridman EA, Beattie BJ, Broft A, Laureys S, Schiff ND. 2014. Regional cerebral metabolic patterns demonstrate the role of anterior forebrain mesocircuit dysfunction in the severely injured brain. Proc Natl Acad Sci U S A 111: 6473–8

Garcia-Borreguero D, Larrosa O, Bravo M. 2003. Parkinson’s disease and sleep. Sleep Med Rev 7: 115–2.

Gatev P, Darbin O, Wichmann T. 2006. Oscillations in the basal ganglia under normal conditions and in movement disorders. Mov Disord 21: 1566–7.

Giacino JT, Whyte J, Bagiella E, Kalmar K, Childs N, et al. 2012. Placebo-controlled trial of amantadine for severe traumatic brain injury. N Engl J Med 366: 819–2.

Gjerstad MD, Wentzel-Larsen T, Aarsland D, Larsen JP. 2007. Insomnia in Parkinson’s disease: frequency and progression over time. J Neurol Neurosurg Psychiatry 78: 476–9

Grabli D, McCairn K, Hirsch EC, Agid Y, Feger J, et al. 2004. Behavioural disorders induced by external globus pallidus dysfunction in primates: I. Behavioural study. Brain 127: 2039–5.

Graham DI, Maxwell WL, Adams JH, Jennett B. 2005. Novel aspects of the neuropathology of the vegetative state after blunt head injury. Prog Brain Res 150: 445–5.

Hannawi Y, Lindquist MA, Caffo BS, Sair HI, Stevens RD. 2015. Resting brain activity in disorders of consciousness: a systematic review and meta-analysis. Neurology 84: 1272–8.

Hazrati LN, Parent A. 1991. Projection from the external pallidum to the reticular thalamic nucleus in the squirrel monkey. Brain Res 550: 142–6

Hutchison WD, Lozano AM, Davis KD, Saintcyr JA, Lang AE, Dostrovsky JO. 1994. Differential Neuronal-Activity in Segments of Globus-Pallidus in Parkinsons-Disease Patients. Neuroreport 5: 1533–3.

Juri C, Chana P, Tapia J, Kunstmann C, Parrao T. 2005. Quetiapine for insomnia in Parkinson disease: results from an open-label trial. Clin Neuropharmacol 28: 185–7

Klein JC, Rushworth MF, Behrens TE, Mackay CE, de Crespigny AJ, et al. 2010. Topography of connections between human prefrontal cortex and mediodorsal thalamus studied with diffusion tractography. Neuroimage 51: 555–6.

Kostopoulos GK. 2001. Involvement of the thalamocortical system in epileptic loss of consciousness. Epilepsia 42 Suppl 3: 13–9

Kreitzer AC. 2009. Physiology and pharmacology of striatal neurons. Annu Rev Neurosci 32: 127–4.

Lant ND, Gonzalez-Lara LE, Owen AM, Fernandez-Espejo D. 2016. Relationship between the anterior forebrain mesocircuit and the default mode network in the structural bases of disorders of consciousness. Neuroimage Clin 10: 27–3.

Laureys S, Faymonville ME, Luxen A, Lamy M, Franck G, Maquet P. 2000. Restoration of thalamocortical connectivity after recovery from persistent vegetative state. Lancet 355: 1790–1

Lazarus M, Huang ZL, Lu J, Urade Y, Chen JF. 2012. How do the basal ganglia regulate sleep-wake behavior? Trends Neurosci 35: 723–3.

Liu X, Lauer KK, Ward BD, Li SJ, Hudetz AG. 2013. Differential effects of deep sedation with propofol on the specific and nonspecific thalamocortical systems: a functional magnetic resonance imaging study. Anesthesiology 118: 59–6.

Llinas R, Ribary U, Contreras D, Pedroarena C. 1998. The neuronal basis for consciousness. Philos Trans R Soc Lond B Biol Sci 353: 1841–9

Lutkenhoff ES, Chiang J, Tshibanda L, Kamau E, Kirsch M, et al. 2015. Thalamic and extrathalamic mechanisms of consciousness after severe brain injury. Ann Neurol 78: 68–7.

Lutkenhoff ES, McArthur DL, Hua X, Thompson PM, Vespa PM, Monti MM. 2013. Thalamic atrophy in antero-medial and dorsal nuclei correlates with six-month outcome after severe brain injury. Neuroimage Clin 3: 396–40.

Magnin M, Morel A, Jeanmonod D. 2000. Single-unit analysis of the pallidum, thalamus and subthalamic nucleus in parkinsonian patients. Neuroscience 96: 549–6.

Mallet N, Pogosyan A, Marton LF, Bolam JP, Brown P, Magill PJ. 2008. Parkinsonian beta oscillations in the external globus pallidus and their relationship with subthalamic nucleus activity. J Neurosci 28: 14245–5.

Mastro KJ, Bouchard RS, Holt HA, Gittis AH. 2014. Transgenic mouse lines subdivide external segment of the globus pallidus (GPe) neurons and reveal distinct GPe output pathways. J Neurosci 34: 2087–9.

McCafferty C, David F, Venzi M, Lorincz ML, Delicata F, et al. 2018. Cortical drive and thalamic feed-forward inhibition control thalamic output synchrony during absence seizures. Nat Neurosci 21: 744–5.

Monti MM. 2012. Cognition in the vegetative state. Annu Rev Clin Psychol 8: 431–5.

Monti MM, Rosenberg M, Finoia P, Kamau E, Pickard JD, Owen AM. 2015. Thalamo-frontal connectivity mediates top-down cognitive functions in disorders of consciousness. Neurology 84: 167–7.

Monti MM, Schnakers C, Korb AS, Bystritsky A, Vespa PM. 2016. Non-Invasive Ultrasonic Thalamic Stimulation in Disorders of Consciousness after Severe Brain Injury: A First-in-Man Report. Brain Stimul 9: 940–4.

Partinen M. 1997. Sleep disorder related to Parkinson’s disease. J Neurol 244: S3–6

Qiu MH, Chen MC, Wu J, Nelson D, Lu J. 2016a. Deep brain stimulation in the globus pallidus externa promotes sleep. Neuroscience 322: 115–2.

Qiu MH, Vetrivelan R, Fuller PM, Lu J. 2010. Basal ganglia control of sleep-wake behavior and cortical activation. Eur J Neurosci 31: 499–50.

Qiu MH, Yao QL, Vetrivelan R, Chen MC, Lu J. 2016b. Nigrostriatal Dopamine Acting on Globus Pallidus Regulates Sleep. Cereb Cortex 26: 1430–9

Saunders A, Oldenburg IA, Berezovskii VK, Johnson CA, Kingery ND, et al. 2015. A direct GABAergic output from the basal ganglia to frontal cortex. Nature 521: 85–9

Schiff ND. 2008. Central thalamic contributions to arousal regulation and neurological disorders of consciousness. Ann N Y Acad Sci 1129: 105–1.

Schiff ND. 2010. Recovery of consciousness after brain injury: a mesocircuit hypothesis. Trends Neurosci 33: 1–9

Schiff ND, Giacino JT, Kalmar K, Victor JD, Baker K, et al. 2007. Behavioural improvements with thalamic stimulation after severe traumatic brain injury. Nature 448: 600–3

Shames JL, Ring H. 2008. Transient reversal of anoxic brain injury-related minimally conscious state after zolpidem administration: a case report. Arch Phys Med Rehabil 89: 386–8

Silva A, Cardoso-Cruz H, Silva F, Galhardo V, Antunes L. 2010. Comparison of anesthetic depth indexes based on thalamocortical local field potentials in rats. Anesthesiology 112: 355–6.

Sutton JA, Clauss RP. 2017. A review of the evidence of zolpidem efficacy in neurological disability after brain damage due to stroke, trauma and hypoxia: A justification of further clinical trials. Brain Inj 31: 1019–2.

Thibaut A, Bruno MA, Ledoux D, Demertzi A, Laureys S. 2014. tDCS in patients with disorders of consciousness: sham-controlled randomized double-blind study. Neurology 82: 1112–8

Tononi G. 2008. Consciousness as integrated information: a provisional manifesto. Biol Bull 215: 216–4.

Trenkwalder C. 1998. Sleep dysfunction in Parkinson’s disease. Clin Neurosci 5: 107–1.

Van Essen DC, Ugurbil K, Auerbach E, Barch D, Behrens TE, et al. 2012. The Human Connectome Project: a data acquisition perspective. Neuroimage 62: 2222–3.

Vetrivelan R, Qiu MH, Chang C, Lu J. 2010. Role of Basal Ganglia in sleep-wake regulation: neural circuitry and clinical significance. Front Neuroanat 4: 145

Visser-Vandewalle V, Temel Y, Ackermans L. 2009. Deep Brain Stimulation in Tourette’s Syndrome In Neuromodulation, pp. 579-86: Elsevier

Whyte J, Rajan R, Rosenbaum A, Katz D, Kalmar K, et al. 2014. Zolpidem and restoration of consciousness. Am J Phys Med Rehabil 93: 101–1.

Williams ST, Conte MM, Goldfine AM, Noirhomme Q, Gosseries O, et al. 2013. Common resting brain dynamics indicate a possible mechanism underlying zolpidem response in severe brain injury. Elife 2: e01157

Xie G, Deschamps A, Backman SB, Fiset P, Chartrand D, et al. 2011. Critical involvement of the thalamus and precuneus during restoration of consciousness with physostigmine in humans during propofol anaesthesia: a positron emission tomography study. Br J Anaesth 106: 548–5.

Xue Y, Han XH, Chen L. 2010. Effects of Pharmacological Block of GABA(A) Receptors on Pallidal Neurons in Normal and Parkinsonian State. Front Cell Neurosci 4: 2

Yoo SS, Kim H, Min BK, Franck E, Park S. 2011. Transcranial focused ultrasound to the thalamus alters anesthesia time in rats. Neuroreport 22: 783–7

Yuan XS, Wang L, Dong H, Qu WM, Yang SR, et al. 2017. Striatal adenosine A2A receptor neurons control active-period sleep via parvalbumin neurons in external globus pallidus. Elife 6

Zheng ZS, Reggente N, Lutkenhoff E, Owen AM, Monti MM. 2017. Disentangling disorders of consciousness: Insights from diffusion tensor imaging and machine learning. Hum Brain Mapp 38: 431–4.

